## Supplemental figure 1 for "Stx2 induces differential gene expression by activating several pathways and disturbs circadian rhythm genes in the proximal tubule"

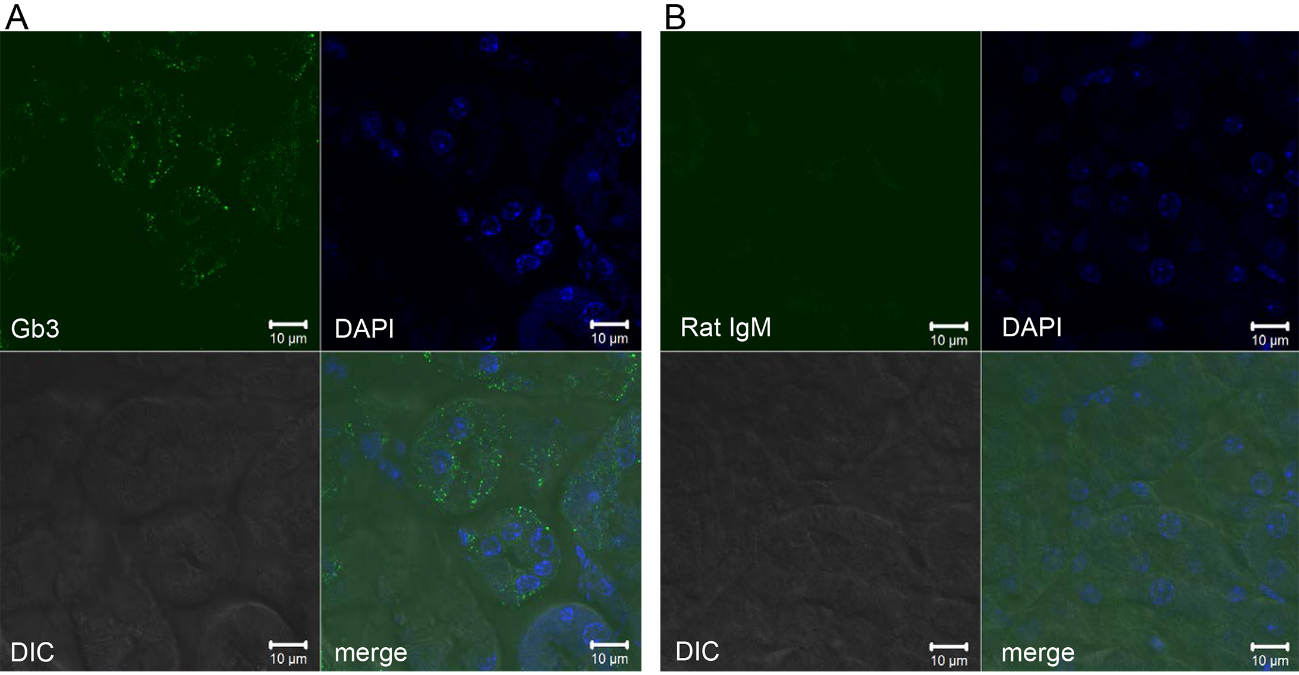

Figure S1. The appearance of isotype control (rat IgM) to Gb3 immunofluorescence staining

(A) Appearance of mouse kidney free-floating section anti-Gb3 immunofluorescence stain with DAPI, DIC and merged pictures. (B) Isotype control (rat IgM) is used in the place of anti-Gb3 antibody to show the negative reaction of normal rat IgM. Rat IgM, DAPI, DIC and merged images.
