## Supplemental figure 2 for "Stx2 induces differential gene expression by activating several pathways and disturbs circadian rhythm genes in the proximal tubule"

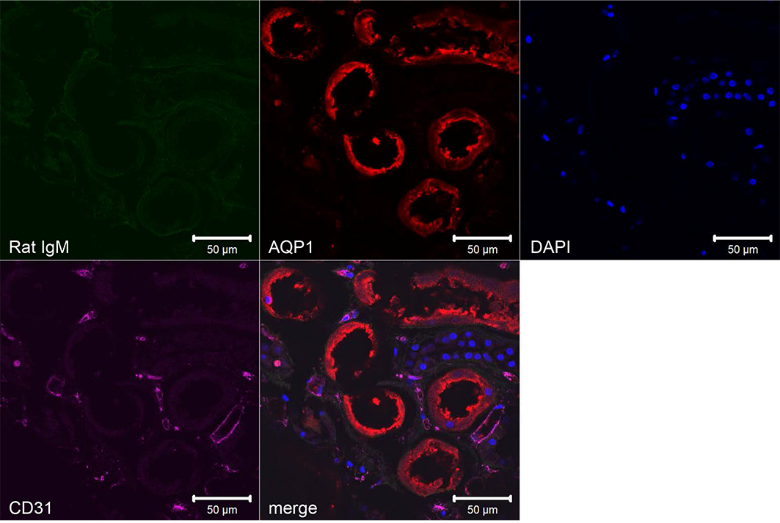

Figure S2. The appearance of isotype control (rat IgM) to Gb3 immunofluorescence staining in human kidney

The appearance of human kidney free-floating section isotype control (rat IgM) is used in the place of anti-Gb3 antibody to show the negative reaction of normal rat IgM. Rat IgM, anti-AQP1, DAPI, anti-CD31 immunofluorescence staining, and merged images.
