## Supplemental figure 3 for "Stx2 induces differential gene expression by activating several pathways and disturbs circadian rhythm genes in the proximal tubule"

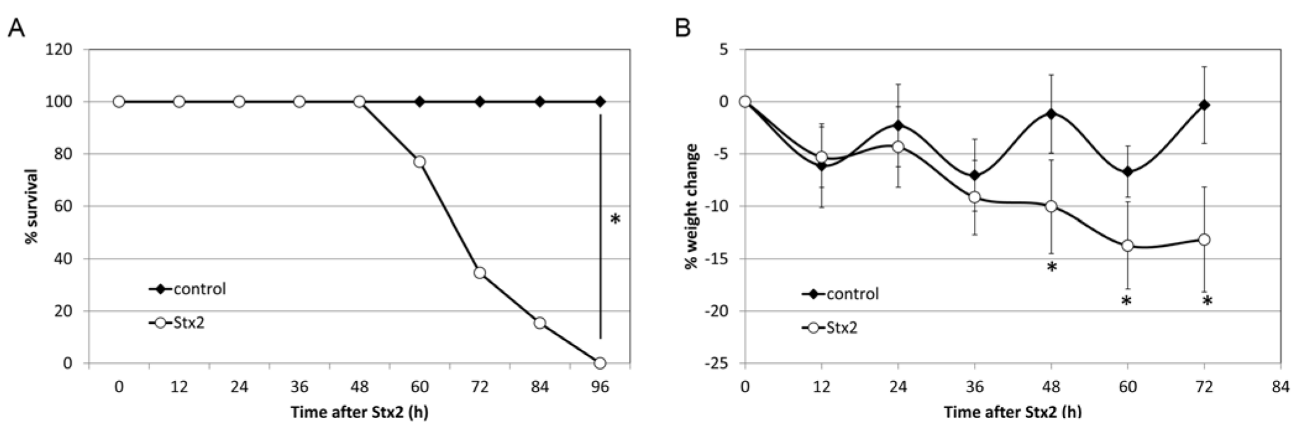
Figure S3. Survival curve and weight changes in saline or Stx2-injected mice. (A) Percent survival of mice injected with saline (closed circle, control) or with Stx2 (open circle, Stx2). The asterisk denotes statistical significance, p<0.05. (B) Percent of weight changes in control or Stx2 mice. Asterisks show p<0.05 between saline control and Stx2.
